# Contribution of mesopelagic fish and cephalopods to the diet of rorquals (*Balaenoptera spp*) and sperm whales (*Physeter macrocephalus*) beyond their feeding grounds

**DOI:** 10.1101/2025.03.14.643344

**Authors:** Cristina Claver, Leire G. de Amézaga, Iñaki Mendibil, Oriol Canals, Rui Prieto, Irma Cascão, Cláudia Oliveira, Mónica A. Silva, Naiara Rodríguez-Ezpeleta

## Abstract

Cetacean conservation requires ecosystem-scale management with special focus on food webs. Rorquals and sperm whales are top predators of complex open ocean food webs and, although mesopelagic fish and cephalopods are predated by these cetaceans, their contribution to their diets is not fully understood. Here, we aimed to better describe the consumption of mesopelagic fish and cephalopods by identifying preferred species consumed by rorquals and sperm whales at mid-latitudes. To do so, we combined the fish and cephalopod community composition inferred from whale faecal and marine environmental DNA samples. We analysed the prey availability and predator preferences by comparing the vertical distribution and abundance of fish and cephalopod species in the water column with the prey items found in faecal samples of rorquals and sperm whales. We found that rorqual consumed mesopelagic fish that perform diel vertical migrations (DVM) such as myctophids. These species were found in depths that matched the deep foraging behaviours during daytime and shallow foraging behaviours during night, confirming that rorquals rely on the DVM to feed at these latitudes. Also, although a high diversity of cephalopods was found across the water column, the faecal content of sperm whales was mainly composed by *Histioteuthis bonellii,* which was abundant between 600 and 1200 meters and matches the diving patterns described for this species in the area. In this study, we present the first comprehensive genetic analysis of the diets of rorquals and sperm whales, expanding our understanding of open ocean trophic ecology to promote effective cetacean conservation.

## Introduction

Rorquals (*Balaenoptera musculus, Balaenoptera physalus, Balaenoptera borealis*) and sperm whales (*Physeter macrocephalus*) are the largest cetaceans and among the most widely distributed species of marine megafauna. Rorquals are filter feeders whose diet relies on small aggregating organisms, such as zooplankton and fish (Aguilar and García-Vernet 2018, Horwood 2018, Sears and Perrin 2018), whereas sperm whales echolocate and directly capture larger individual preys, such as cephalopods (Whitehead 2018). These large predators, characterized by their high mobility and energy demands, require vast, healthy habitats and abundant food resources to sustain their massive body sizes (Williams 2006). Indirect threats such as prey decline and range shifts, driven by climate change and overfishing, are among the often overlooked impacts they face (Bearzi and Reeves 2021). Therefore, effective conservation of these species depends on the preservation of entire ecosystems, including protecting food webs within their habitats. Because achieving a global sustainable management strategy is challenging (Mason, Ward et al. 2020), as an alternative feasible approach, prioritizing critical areas of preservation by identifying feeding grounds and preferred preys has been proposed (Stephenson, Hewitt et al. 2021).

Mesopelagic fish and cephalopods play a crucial role in food webs, connecting deep-sea and surface ecosystems through their diel vertical migrations (DVMs) (Irigoien, Klevjer et al. 2014, Judkins and Vecchione 2020). Their high biomass has attracted interest from fishing industries (FAO 2014), raising concerns about the impact of their exploitation on the complex and highly interconnected open ocean ecosystem. Rorquals and sperm whales are known consumers of these taxa (Pauly, Trites et al. 1998, Cook, Bernard et al. 2020) and diel foraging patterns related to DVMs of mesopelagic preys such as krill or cephalopods have been reported in these cetaceans (Croll, Tershy et al. 1998, Goldbogen, Calambokidis et al. 2006, Aoki, Amano et al. 2007), suggesting a high dependence on a mesopelagic-based diet. However, identifying their targeted preys is challenging either through direct observations, due to foraging occurring far from the surface, or through stable isotope analysis, given the limited taxonomic resolution of the method. Research in rorquals has mainly focused of understanding the consumption of zooplankton and krill, while sperm whale diet studies are limited to cephalopods, with little effort dedicated to understanding the contribution of other prey items.

The Azores archipelago offers essential foraging grounds and breeding areas that are crucial for the complex life cycles of rorquals and sperm whales. Located in the Mid-Atlantic Ridge, these islands feature a complex topography that includes abundant seamounts and deep bathymetries (Peran, Pham et al. 2016). These physiographic features, combined with the regiońs unique oceanographic conditions and processes, make the Azores a marine biodiversity hotspot and a significant biogeographic area for large migratory whales (Silva, Prieto et al. 2013, Prieto, Silva et al. 2014). Rorquals pause their northward migration in Azorean waters seasonally, suggesting they stop there to forage (Silva, Prieto et al. 2013). Male sperm whales also migrate to the Azores from northern latitudes during summer to mate, while females and juveniles remain in these waters throughout the year (Steiner, Lamoni et al. 2012). This indicates a consistent and significant prey availability (Silva, Prieto et al. 2014). In the Azores, fin whales (*Balaenoptera physalus*) adjust their diel foraging activity by diving deeper (∼100 meters, down to 300m) during the day and shallower (∼50 m) at night (Fonseca, Pérez-Jorge et al. 2022). Isotopic analyses reveal that prey of higher trophic levels than zooplankton significantly contribute to the diet of fin whales in the region (Silva, Borrell et al. 2019). These findings combined (Woods and Barkmann 1995) suggest that fin whales may target mesopelagic nekton, such as fish. In contrast, sperm whales consistently forage between 600 and 1000 m (Oliveira, Pérez-Jorge et al. 2022), indicating they may focus on deep-water cephalopod prey species.

Recent studies have provided some insights into diel foraging patterns of rorquals and sperm whales (see references above), but their diets remain poorly understood due to challenges in obtaining samples, limited information on their deep-sea prey species, and variability in prey availability across their distribution ranges. Much of what is known about their diets dates back to the whaling era through visual analysis of stomach contents (Sergeant 1966, Roe 1969, Martin and Clarke 1986, Flinn, Trites et al. 2002). Yet, with the global ban on commercial whaling, stomach content analysis is now only possible on stranded individuals, which do not accurately represent healthy populations and are rare. For example, over the last decade, only two studies have examined stomach contents of stranded animals, including six sperm whales (Tonay, Öztürk et al. 2021, Similä, Haug et al. 2022) and one fin whale, the latter with an empty stomach (San Martín, Paso Viola et al. 2021).

Dietary DNA analysis, which consists on the study of DNA from stomach contents or faeces, is becoming popular in in marine trophic ecology studies (de Sousa, Silva et al. 2019) for its accuracy and non-invasiveness. Due to the limitations in the analysis of stomach contents, whale faeces represent a critical resource as records of recently ingested preys for understanding top-predator diets (Arregui, Borrell et al. 2018). Although still relatively rare, genetic studies of cetacean faeces have already enhanced our understanding of prey contributions (Ford, Hempelmann et al. 2016) and revealed previously unknown or understudied prey items (De Vos, Faux et al. 2018, Carroll, Gallego et al. 2019). Another emerging approach is the analysis of environmental DNA (eDNA), i.e., DNA released by macrofauna to the environment, which combined with other methodologies, are enabling the study of cetacean interactions with prey such as cephalopods (Visser, Merten et al. 2021) and fish (Berger, Bougas et al. 2020). Together, these molecular approaches can update dietary research, providing a precise and comprehensive view of contemporary cetacean feeding habits and ecological roles.

The main objectives of this study are 1) to better understand the consumption of mesopelagic fish and cephalopods by rorquals and sperm whales, and 2) to determine if these predators exhibited prey preferences. To do so, we analysed the prey availability and predator preferences by comparing the vertical distribution and relative abundance of fish and cephalopod species in the water column within the diving range of rorquals and sperm whales with the prey found in their faecal samples. This study provides the first detailed genetic analysis of sperm whale and rorqual diets contributing to a better understanding of prey preferences and the role of mesopelagic resources to these top predators. Such knowledge is crucial to understand trophic interactions and dynamics in open ocean ecosystems and to provide insights into impacts of food-web perturbations.

## Material and Methods

Complementary Material and Methods information can be found in Appendix 2 of Supplementary Material. Raw sequencing reads and associated metadata are available on the NCBI SRA (BioProject XX). Developed scripts and corresponding output files are available at GitHub (https://github.com/cclaver001/azores_v2).

### Sample collection and DNA extraction

Whale faeces were collected opportunistically between 2011 and 2021 around Faial and Pico islands in the Azores archipelago, Portugal (**Figure 1**). A total of 34 samples from three rorqual species (*Balaenoptera musculus, B. physalus, B. borealis*) and sperm whales (*P. macrocephalus*) were included in this study (**Table S1**). Samples were taken using a nylon stocking secured over a metal ring attached to a pole. Chunks were randomly sampled for each faeces (**Figure S1**). Samples were stored in a cooler onboard until landing and subsequently frozen at -80°C until DNA extraction. Water samples were collected in the same area during June 2021 onboard the OceanXplorer research vessel. Six vertical profiles were sampled twice, once during day and once during night with Niskin bottles mounted on a CTD rosette. A total of 66 two-liter samples were taken in triplicate between 10 and 1200 m (**Table S2**). Additionally, Milli-Q water was filtered onboard in triplicate and used as negative control. Water samples were filtered through the 47 mm GN Metricel® MCE Membrane Disc Filters with a 0.45 µm pore size and stored frozen.

**Figure 1.**
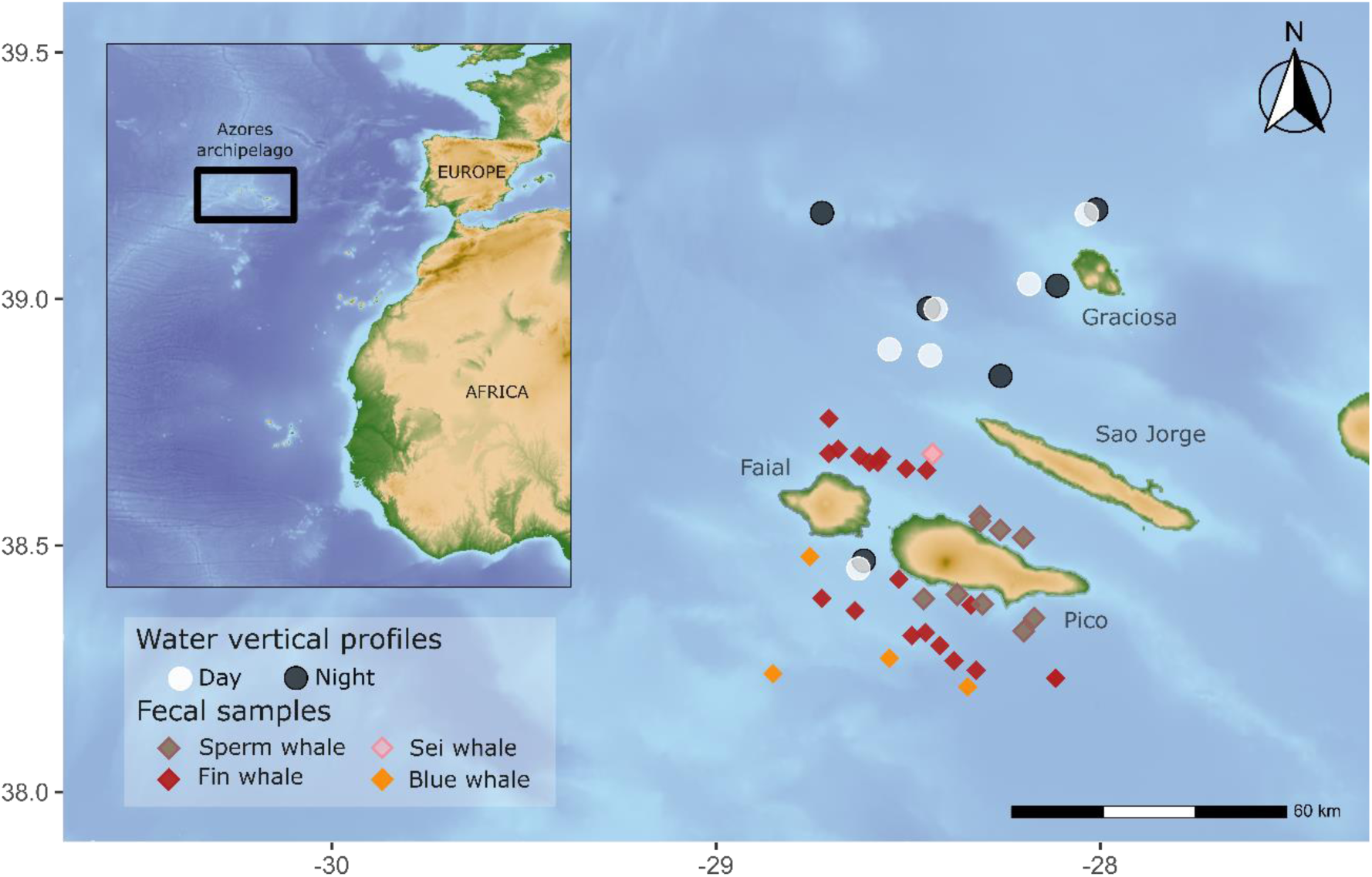
Location of water vertical profile sampling stations (white and black) and cetacean faeces (diamonds coloured per species).

Faecal DNA was extracted using the QIAmp Fast Stool Mini Kit (Qiagen) following manufacturer’s instructions. An additional cleaning step was performed with DNeasy PowerClean Pro Cleanup Kit (Qiagen) according to the manufacturer’s protocol. Water DNA was extracted from the filters using the DNeasy PowerWater Kit (Qiagen) following manufacturer’s protocol. DNA concentration (µg/ml) was calculated with fluorimetry (Qubit), and sample cleanliness was determined with UV spectrometry (NanoDrop). In addition, DNA quality was assessed by electrophoresis, migrating DNA on a 1.0% agarose SYBR safe (Thermo Fisher) TAE buffer gel. Extracted DNA samples were eluted to 10 ng/µl. DNA extracts were stored at -20°C until further analysis.

### Library preparation and sequencing

Libraries were generated following a modified version of the *tagsteady* protocol (Carøe and Bohmann 2020) targeting two barcodes. The first primer set was designed for fish and targeted the 12S rRNA gene, yielding a ∼63 base pair amplicon (*teleo*_forward 5’ACACCGCCCGTCACTCT3’ and *teleo*_reverse 5’CTTCCGGTACACTTACCATG3’) (Valentini, Taberlet et al. 2016). The second primer set was designed for amplifying cephalopods and targeted a 140-190 base pair region of the 18S rRNA gene (Ceph18S_forward 5’CGCGGCGCTACATATTAGA3’ and Ceph18S_reverse, 5’GCACTTAACCGACCGTCGAC3’) (De Jonge, Merten et al. 2021). Because the *teleo* primers coamplify nontarget species that are abundant in the samples, two blocking primers were used: a human blocking primer for water samples (*teleo*_blk ACCCTCCTCAAGTATACTTCAAAGGAC- SPC3I) (Valentini, Taberlet et al. 2016) and a whale blocking primer developed in this study for the feces (teleo_cetaceaBP CCCGTCACCCTCCTCAAGTA). Briefly, a cetacean-exclusive sequence overlapping the *teleo* forward primer was identified and the blocking oligo was developed following recommendations in Vestheim, Deagle et al. (2011). A detailed description of the whale blocking primer design can be found in Supplementary Material (Appendix 2).

Each DNA sample was amplified in triplicate using a different set of tagged primers including 6 bases for tag, one or two degenerated bases (Ns) and the forward and reverse *teleo or ceph18S* primer pair, in a total volume of 15 µl using 7.5 µl of Phusion Master Mix with HF Buffer (Thermo Fisher); 3 µl of the primer [0,2μM], 3 µl of DNA [10 ng], and 1.5 µl of ddH2O. For *teleo*, 0.3 µl of blocking primer [2μM] were also included and ddH2O was reduced to 1.2 µl. The PCR conditions were as follows: pre-denaturation at 98°C for 3 mins; 40 and 35 cycles denaturation at 98°C for 10 seconds for *teleo* and *ceph18S*, respectively; annealing at 55° and 62° for 30 seconds for *teleo* and *ceph18S*, respectively; extension at 72°C for 45 s; and a final extension at 72°C for 10 min. PCR negative controls were included for each pool of samples, resulting in 7 and 6 controls for *teleo* and *ceph18S*, respectively. Also, positive controls (DNA extracted cat fur) were included to monitor tag-jumping (Rodriguez-Martinez, Klaminder et al. 2023) or index hopping (van der Valk, Vezzi et al. 2020). Amplification was visually confirmed on an electrophoresis 1.2% agarose TBE buffer gel and positive replicate PCR products were mixed at equimolar ratios based on gel bands strength. The mixes were then purified using AMPure XP beads (Beckman Coulter, California, USA) following the manufacturer’s instructions, and Illumina’s compatible adapters were ligated to each mix as in Carøe and Bohmann (2020). The mixes were purified with AxyPrep MAG PCR Clean-Up Kit (Axygen™), end-repaired and adjusted to 30 μl with molecular water and mixed, afterwards, with 10 μl of end-repair master mix each. Libraries were purified with 100 µl AxyPrep MAG PCR Clean-Up Kit (Axygen™) and measured with Qubit HS kit before mixing the different pools equimolarly. Sequencing was performed on the Illumina NovaSeq using NovaSeq SP 150 PE cycle.

### Read pre-processing, taxonomic assignment and quality control

Raw reads were demultiplexed using cutadapt (Martin 2011) based on tags allowing for no error rate (-e 0), and quality was verified using FASTQC (Andrews 2010). Forward and reverse reads were merged using PEAR (Zhang, Kobert et al. 2014) with a minimum overlap of 20 and 10bp for *teleo* and *ceph18S*, respectively. Trimmomatic (Bolger, Lohse et al. 2014) was used to retain assembled sequences with a minimum average Phred score of 25 for subsequent analyses. Forward and reverse primers were removed with cutadapt (Martin 2011) and only reads containing both forward and reverse primers were kept, allowing a maximum error rate of 20%. Reads longer than 50 or 100 nucleotides were retained for *teleo* and *ceph18S* amplicons, respectively. Because forward and reverse sequences are expected in both R1 and R2 files, the combinations of forward/reverse complement primers and reverse/forward complement primers were searched in parallel and later sequences were joined. For *teleo,* sequences were aligned against a 12S reference database (covering the *teleo* region) to discard amplicons not covering the target region. Using *mothur* (Schloss, Westcott et al. 2009), sequences with ambiguous bases were discarded and potential chimeras were identified based on the *UCHIME* algorithm (Edgar, Haas et al. 2011) and removed. Samples with less than 1,000 reads were discarded.

For each barcode, a local reference database was created for taxonomic assignment. First, all sequence records corresponding to 12S and 18S rRNA genes from North Atlantic vertebrates and invertebrates respectively were retrieved from the MetaZooGene Atlas and Database (MZGdb). Second, from the 12S and 18S retrieved reads, only those belonging to respectively fish (filtered by “*Saltwater*” and species classified as “stray/questionable occurrence” omitted) and cephalopod species inhabiting the Azorean archipelago were selected using the “*Information by Country*” tool provided by Fishbase and SeaLifebase. Taxonomic assignment was performed for each unique sequences using the naïve Bayesian classifier method (Wang, Garrity et al. 2007) implemented in *mothur.* The confidence threshold for accepting the classifications was set to 70% for both barcodes to provide stringent results. Reads assigned to the same species, or to the same genus or family (when respectively species or genus level classification was not possible) were grouped into phylotypes. Only phylotypes assigned to family, genus or species level were kept for subsequent analysis. Phylotypes with less than 10 reads per sample were discarded (**Tables S3-8**).

All sampling and laboratory negative controls resulted in very few and unclassified reads, except for the sampling control amplified with *ceph18S*, which resulted in a high number of *Loligo forbesii* reads, probably resulting of onboard contamination with this species. For the *teleo* dataset, an additional taxonomic assignment was performed using an enlarged database including mammals, allowing to confirm that no sequences from the positive control (i.e., cat DNA) were found in the samples. In all faecal samples reads corresponding to the predator (mean ≈ 12% of the reads) were amplified and sequenced. For two fin whale samples, whale sequences were assigned to blue whale (sample BPH_58) and sei whale (BPH_146).

### Data analysis and statistical tests

Water samples were divided into four depth layers: upper epipelagic (<100 m), lower epipelagic (100-300 m), upper mesopelagic (300-600m) and lower mesopelagic (600-1200m). These categories correspond to the depth ranges where fin whales forage during night and day (Fonseca, Pérez-Jorge et al. 2022), the location of the deep scattering layer in Azores (Cascao, Domokos et al. 2019) and the foraging depths of sperm whales (Oliveira, Wahlberg et al. 2016), respectively. Water samples were merged by the described depth layers and relative read abundance (RRA) was calculated for each depth range category for day- and night-time collected samples. For faeces, nonmetric multidimensional scaling (NMDS) (*vegan* package) based on Bray-Curtis distances was applied to visualise the similarity of sample composition, and the ANOSIM test was used to determine statistical significance of the grouping of samples from different predators. The contribution of phylotypes to the ordination of samples was calculated using *envfit* function (vegan package) and the phylotypes that were contributing most significantly (R ≥ 0.7; p<0.05) were plotted. Overall, RRA was calculated for rorquals (sei, blue and fin whales combined) and sperm whales, as well as for each individual sample. Frequency of prey occurrence was calculated for sperm whale and rorquals. A given prey was considered to be present if it accounted for more than 0.1% of the reads of that sample. This threshold was established following considerations discussed in Deagle, Thomas et al. (2019) to avoid secondary consumption and environmental contaminations. Only samples with more than 1,000 fish or cephalopod reads were analysed individually. All statistical analyses were conducted in RStudio (R version 4.2.2; R Core Development Team).

## Results

### Fish and cephalopod diversity, occurrence and relative abundance in water and faecal samples

Overall, 23 fish phylotypes (17, 4 and 2 classified respectively to species, genus and family level) and 27 cephalopod phylotypes (22, 1 and 4 classified respectively to species, genus and family level) were found in water and faecal samples (**Figure 2 a,b**). The highest number of both fishes and cephalopods was found in water samples. Considering relative abundance, fish community composition was similar in water and rorqual faecal samples, while sperm whale faeces were mainly composed by the deep sea *Cryptopsaras couesii* and Myctophidae (**Figure 2c**). Cephalopod community composition was also diverse in water and rorqual faecal samples, whereas it was dominated by *Histioteuthis bonellii* in sperm whale faeces (**Figure 2d**).

**Figure 2.**
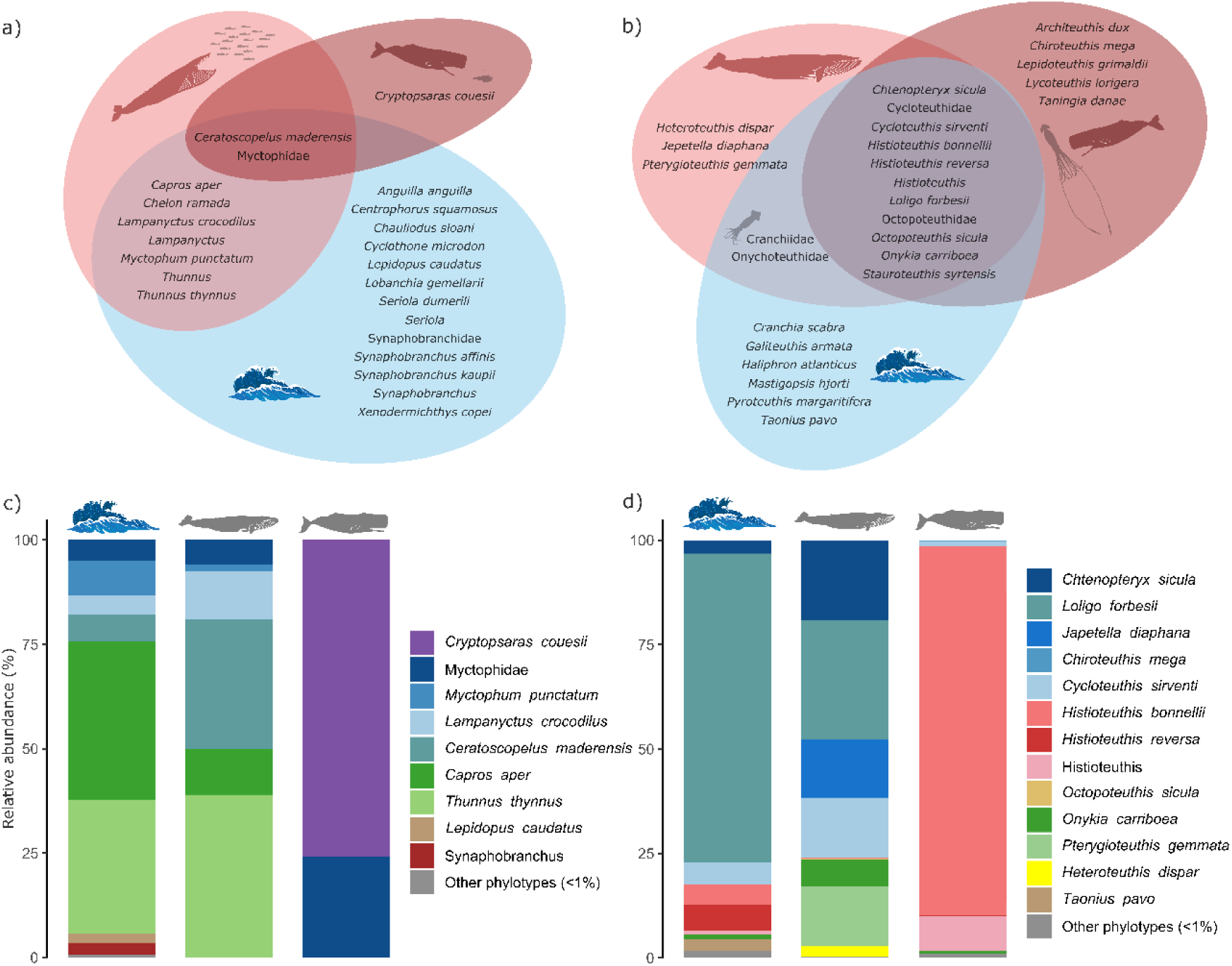
Differences and similarities across water, rorqual faeces and sperm whale faeces samples. Venn diagrams represent shared taxa for each group: **a)** fishes and **b)** cephalopods, and bar plots represent relative abundance (%) of **c)** fish and **d)** cephalopod.

Sperm whale and rorqual diets show statistically significant differences for both fish (R=0.34) and cephalopods (R=0.81) (**Figure 3**). Despite these differences, no fish species was clearly driving the separation between sperm whale and rorqual diets; however, for cephalopods, several groups such as *Cycloteuthis sirventi* and histioteuthids were primarily responsible for the separation. For both rorquals and sperm whales, the number of fish prey in each faecal sample was small (<6) (**Figure 4a**). The fish species with the highest occurrence in rorqual faecal samples was *Ceratoscopelus maderensis*, yet it was only present in one third of the samples (**Figure 4b**) and, in sperm whale samples *C. couesii*, present in half of the samples (**Figure 4c**). The number of cephalopod preys per individual was substantially higher in sperm whale samples than in rorqual samples, except for two samples (**Figure 4d**). In rorqual samples, each cephalopod prey was present in a small percentage of samples (<12%, corresponding to less than 3 samples), with *L. forbesii* being present in almost one quarter of the samples **(Figure 4e)**. In contrast, in sperm whale samples histioteuthids were present in all samples and *Onykia carriboea* and *C. sirventi* were present in 85% of the samples **(Figure 4f)**.

**Figure 3.**
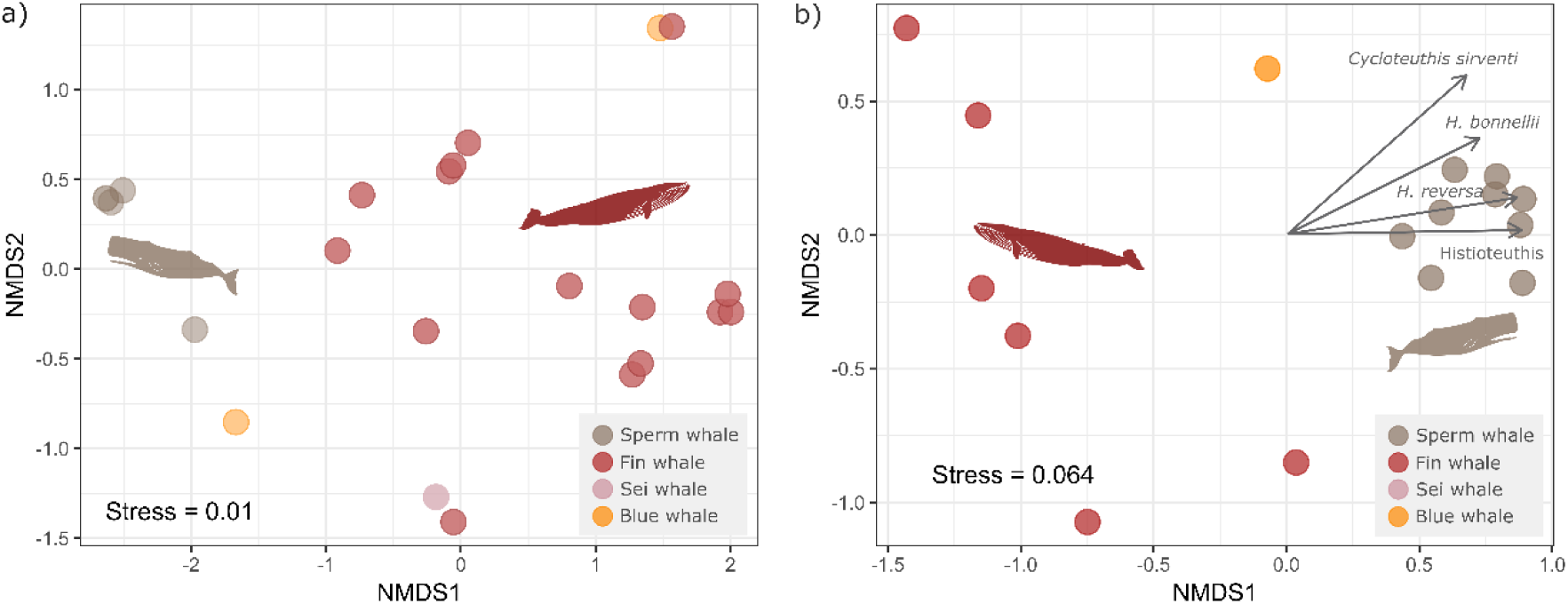
Non-metric multidimensional scaling (NMDS) of rorqual and sperm whale faeces for **a)** fishes (ANOSIM test: R=0.34; p<0.05) and **b)** cephalopods (ANOSIM test: R=0.81; p<0.05).

**Figure 4.**
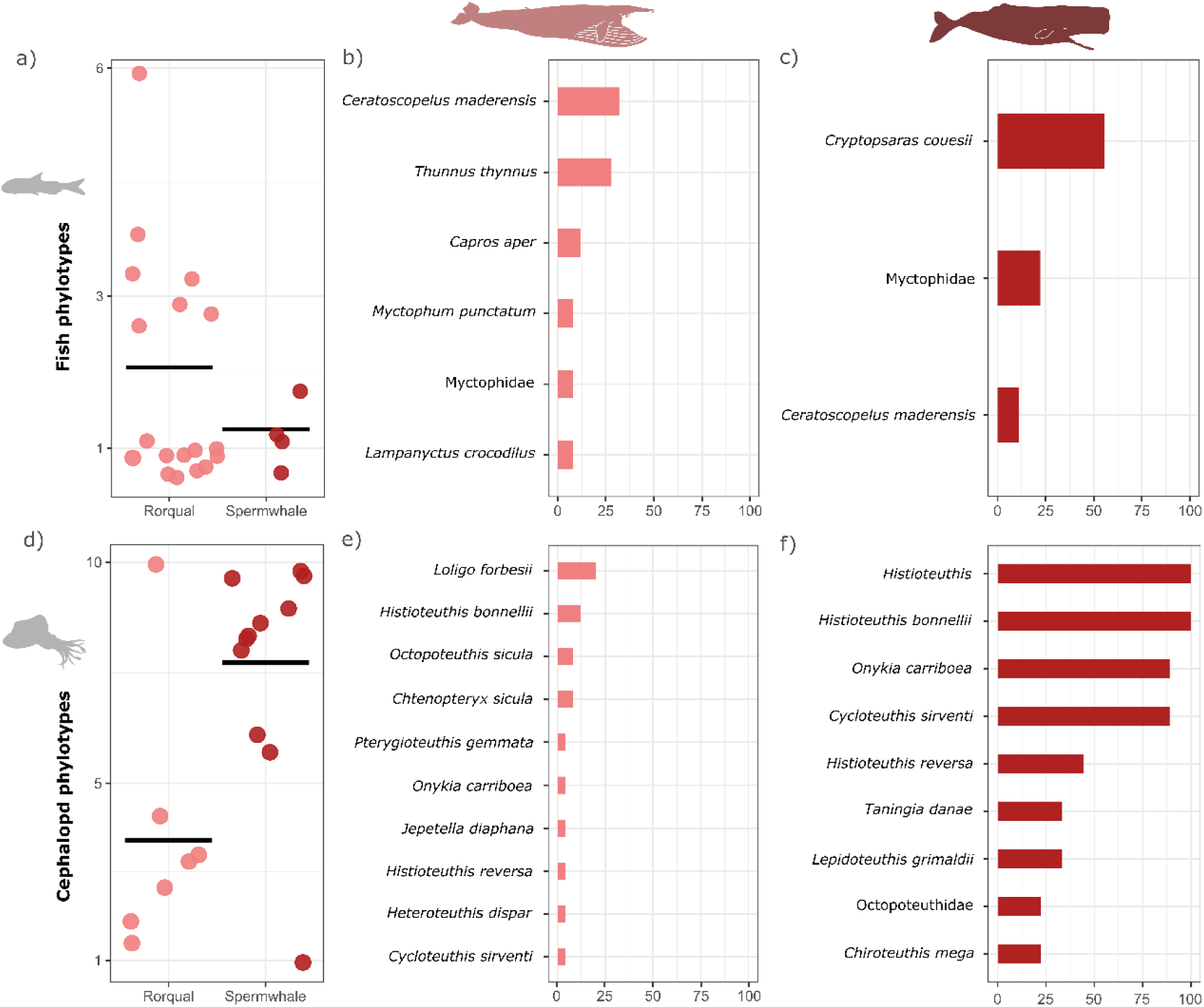
Contribution of different prey to cetaceans diets. **a)** Fish richness (number of phylotypes) in rorqual (left; light red coloured dots) and sperm whale (right; dark red coloured dots) faecal samples. Frequency of occurrence (% of samples) of fish phylotypes in **b)** rorqual and **c)** sperm whale faecal samples. **d)** Cephalopod richness (number of phylotypes) in rorqual (left; light red coloured dots) and sperm whale (right; dark red coloured dots) faecal samples. Frequency of occurrence (%) of cephalopod phylotypes in **e)** rorqual and **f)** sperm whale faecal samples.

### Prey availability in depth-layer samples and composition of faecal samples

No fish DNA was detected in 7 out of 25 (30%) rorqual and 5 out of 9 (55%) sperm whale faeces (**Figure 5a**). In the sei whale sample, *Lampanyctus crocodilus* was found, which was also present in two fin whale samples. Of the four blue whale samples, only two had fish DNA and corresponded to Myctophidae and *Capros aper*. In water samples, the highest fish diversity was found in the upper mesopelagic zone, although some species such as *Chauliodus sloani*, *Cyclothone microdon* and *Xenodermichthys copei* were exclusively found in the lower mesopelagic zone (**Figure S2a**). During the day, the upper epipelagic zone was mainly composed of *Thunnus thynnus*, whereas myctophids were mostly found in the upper mesopelagic layer. At night, the upper epipelagic zone was dominated by *C. aper* and myctophids were distributed throughout the water column, with *C. maderensis* found mainly in upper epipelagic waters **(Figure 5b)**.

**Figure 5.**
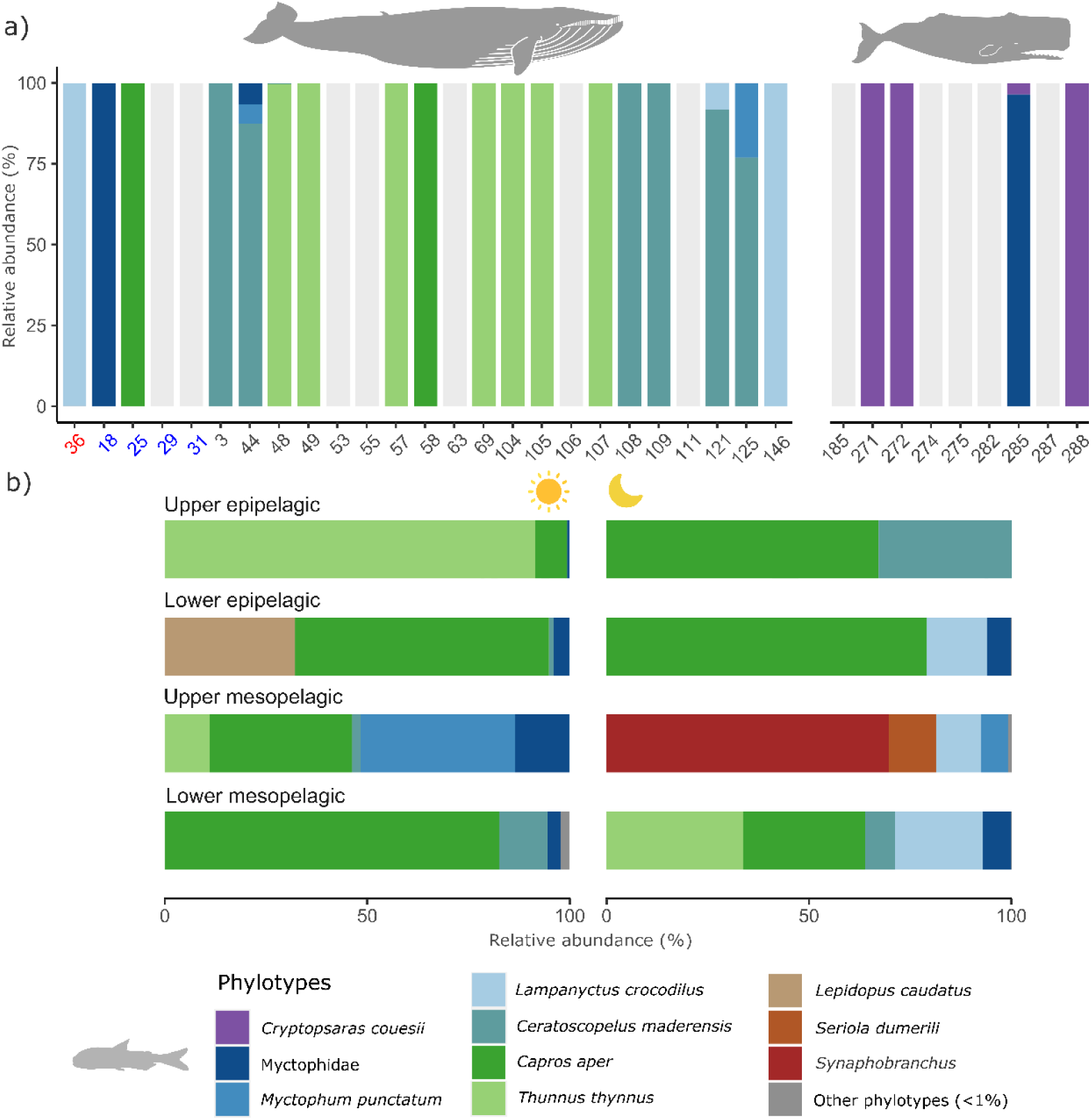
Relative fish abundance in **a)** faecal samples and **b)** across the water column during day (left) and night (right). Greyish bars correspond to empty samples. Rorqual faeces with their name in red corresponded to sei whale (36) and blue corresponded to blue whales (18, 25, 29, 31) while the rest of samples (3 to 146) corresponded to fin whales and sperm whales (185-288).

Regarding cephalopods, 18 out of 25 (70%) rorqual samples were empty, while all the sperm whale samples had cephalopod DNA, which were mainly composed by *H. bonnellii* (**Figure 6a**). The sei whale sample was empty and only one blue whale sample had cephalopod DNA corresponding to *C. sirventi*. The whole water column was mainly composed by *L. forbesii,* and the lower mesopelagic zone presented the highest diversity (**Figure S2b**). *O. carriboea* was found in the upper epipelagic and histioteuthids were abundant in the mesopelagic zone. Samples corresponding to the lower epipelagic zone at night had no cephalopod reads and no evidence of DVM was found **(Figure 6b).**

**Figure 6.**
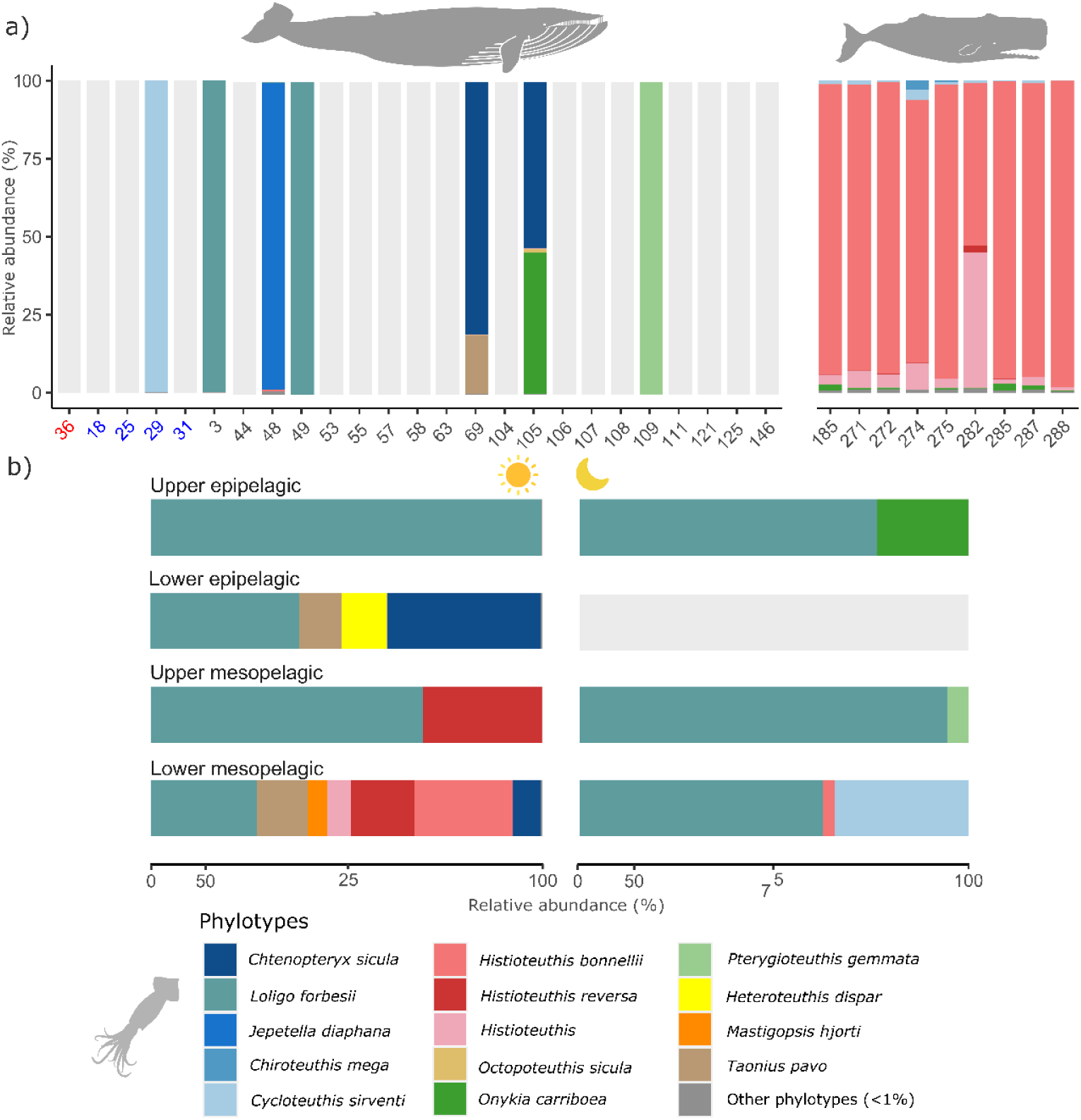
Relative cephalopod abundance in **a)** faecal samples and **b)** across the water column during day (left) and night (right). Greyish bars correspond to empty samples. Rorqual faeces with their name in red corresponded to sei whale (36) and blue corresponded to blue whales (18, 25, 29, 31) while the rest of samples (3 to 146) corresponded to fin whales and sperm whales (185-288). For water samples, the lower epipelagic zone at night, which had no cephalopod DNA.

## Discussion

This research is pioneering in utilizing multi-source DNA to elucidate predator-prey interactions and reveals the high potential of eDNA for studies on trophic ecology of large pelagic whales. In one hand, opportunistic cetacean faeces collection provides valuable trophic information through DNA analysis, while on the other hand, water samples reveal the richness and vertical distribution of cephalopod and fish communities in the area, allowing to examine trophic interactions based on prey availability. Notably, while our study does not allow to quantify the relative contribution of fish and cephalopods to the overall rorqual and sperm whale diet, it provides novel information regarding these two taxonomic groups as cetacean prey.

### Rorquals feed on both epipelagic and mesopelagic fish

Although the contribution of fish to the diets of some toothed whales is significant (Ford and Ellis 2006, Nielsen, Teilmann et al. 2019, Iglesias, Santora et al. 2023), the fish prey of large migratory baleen whales are not well-documented, with most research focused on stable isotope analysis (Witteveen and Wynne 2016) and limited to specific species such as humpback whales (*Megaptera novaeangliae*) (Witteveen, Worthy et al. 2012, Fleming, Clark et al. 2016, Witteveen and Wynne 2016). North Atlantic fin and sei whales are known to consume small schooling fish in their high-latitude foraging grounds (Christensen, Haug et al. 1992, Sigurjónsson and Víkingsson 1997), despite this being primarily described in higher latitudes. Our results show that 76% of rorqual faeces had fish DNA, indicating that North Atlantic rorquals consume fish. Myctophids were predominant in the faeces of all three rorqual species, together with *C. aper*. The data demonstrate diel variations in the distribution of *C. aper* within the water column, showing higher abundance in the upper mesopelagic zone during nighttime compared to daytime, which aligns with the reported DVMs of this species in the Azores (Cascao, Domokos et al. 2019). Similarly, myctophids (i.e., *C. maderensis, L. crocodilus* and Myctophidae) were primarily detected at depths between 300 and 600 meters during the day, consistent with the depth range of the deep scattering layer in the Azores (Cascao, Domokos et al. 2019), but exhibited increased abundance in the epipelagic zone at night. These observations suggest that rorquals capitalize on the elevated availability of *C. aper* and myctophids in shallower waters during nighttime. Nevertheless, due to the limited sample size for sei and blue whales, these results should be interpreted with caution.

The detection of *Thunnus thynnus* in rorqual faeces was unexpected. Although this species is common in the Azores, especially in spring when rorquals are present in the area (Arrizabalaga, Pereira et al. 2008, Arregui, Galuardi et al. 2018), it is highly unlikely that rorquals prey on adult and juvenile bluefin tunas. Furthermore, bluefin tuna do not spawn in the Azores, so these detections could not refer to eggs or larvae in the ichthyoplankton. An explanation for the presence of tuna DNA in rorquals could be that tuna DNA was very abundant in the water filtered by rorquals when feeding. Yet, it has been reported that tuna aggregate with other megafauna such as whale sharks (Fontes, McGinty et al. 2020) or dolphins (Clua and Grosvalet 2001) to hunt. We cannot totally rule out a contamination, but the fact that this species does not appear in the extraction and PCR negative controls suggests that a biological explanation is likely.

### Beyond common perceptions: the overlooked prey of rorquals

Rorqual whales are often considered specialized predators, with little variety of prey and clear preference for zooplankton taxa such as euphausiids or calanoid copepods (Aguilar and García-Vernet 2018, Horwood 2018, Sears and Perrin 2018) but recent research is broadening the prey spectrum of these whales (e.g. De Vos, Faux et al. 2018). Here, we found a variety of cephalopod phylotypes in 7 rorqual faeces. Indeed, cephalopod intakes have been described for sei, fin and blue whales, although it has been considered an indirect ingestion while preying upon other preys (Kawamura 1980, Clarke 1996). This might be the case of *Stauroteuthis syrtensis,* a deep-sea octopus that was found in rorqual faeces. This species mainly feeds on calanoid (Jacoby, Youngbluth et al. 2009), which are common prey of rorquals. Thus, *S. syrtensis* might have been inadvertently swallowed by whales during foraging on copepods. Likewise, some of the cephalopods found in rorqual samples are part of the macroscopic zooplankton communities, and they can form mixed shoals with other organisms. For instance, *Heteroteuthis dispar* was found in fin whale samples and species in this genus form mixed schools with shrimps (Jereb and Roper 2005); so, they can be ingested together due to lack of selectivity when whales target patchy preys (De Vos, Faux et al. 2018). Although some of the cephalopod species found in rorqual samples are bathypelagic, such as *Japetella diaphana*, the juveniles of these species are found in the epipelagic zone (Judkins and Vecchione 2020) where they can be hunted together with other zooplankton taxa. Others, such as *Pterygioteuthis gemmata* are small (∼3 cm) mesopelagic species that perform DVMs and rise to shallow waters at night (Guerra, González et al. 2014), where they become accessible to rorqual whales.

Rorquals may also directly target cephalopod aggregations. The most frequent species in rorqual faeces was *L. forbesii*, which was abundant in our samples and forms schools in the area (Pierce, Sauer et al. 2013, Guerra, González et al. 2014). Overall, all the cephalopod species found in rorqual samples are schooling, small or part of the zooplankton community in early stages. However, the low occurrences and variability of cephalopod phylotypes among samples suggests that these preys are not frequent, and more samples would be needed to have a representative dataset with more consistency between faecal contents. Whether or not rorquals aim to capture them, cephalopod consumption may be a supplementary source of nutrients during migrations in the absence of other prey. Very little research has been done in cephalopod contribution to baleen whale diets and this study represents the most detailed work in this field.

### Sperm whales exhibit prey preferences within their foraging depths

About 30 mesopelagic cephalopod species have been identified as sperm whale prey in the Azores (Silva, Fonseca et al. 2022). The families Histioteutidae and Octopoteuthidae are the primary contributors to prey mass (∼70%) (Clarke, Martins et al. 1993), with *H. bonnellii* alone accounting for 60% of the specimens (Clarke 1956). In our samples, *Histioteuthis spp.* (including *H. bonellii*, *H. reversa* and unidentified *Histioteuthis* species) represented >80% of the DNA sequences and were present in all sperm whale faeces, whereas octopoteuthids (*Octopoteuthidae*) were found in one quarter of the samples with low abundance (<1%). Other known sperm whale prey identified in our samples included *Architeuthis dux, Taningia danae, Lepidoteuthis grimaldii*, *C. sirventi* and *Chiroteuthis mega* (Guerra, González et al. 2014, Silva, Fonseca et al. 2022), although they were present in low abundance and frequency.

Our study identified *H. bonnellii* as the predominant cephalopod species in the faeces of sperm whales. This species is notably abundant in the lower mesopelagic zone (600-1200 m), which aligns with the documented diving patterns of sperm whales in this region (Oliveira 2014, Oliveira, Pérez-Jorge et al. 2022). Interestingly, during daytime, this depth layer exhibited high cephalopod diversity, including other abundant phylotypes such as *L. forbesii* and *Taonius pavo*, thought these species were either absent or minimally represented in the whale faeces. Furthermore, at night, the relative abundance of *H. bonnellii* significantly decreased, while other species, especially *L. forbesii* become dominant in the lower mesopelagic zone. Sperm whales forage both day and night within the same depth ranges (Oliveira 2014, Oliveira, Pérez-Jorge et al. 2022), indicating a selective predation pattern that favours histioteuthids, despite the availability of other species. Clarke, Martins et al. (1993) documented that sperm whales primarily consume slow-swimming, luminous, neutrally buoyant squids such as *Histioteuthis spp,* rather than by pursuing faster swimming, larger cephalopods. Our findings corroborate these results and strongly suggest that prey vulnerability may exert a greater influence on sperm whale feeding behaviour than prey availability. These insights contribute to a deeper understanding of sperm whale-prey dynamics and underscore the potential impacts of shifts in in prey distribution and abundance on sperm whale populations.

Besides the species that coincide between the literature and our findings, there are other less described prey species, such as *O. carriboea,* which constitutes the third most common prey in faeces, although in relatively small proportion. Notably, this species was only detected in night samples from upper epipelagic waters. Juveniles of *O. carriboea* have been reported to occur in the epipelagic zone and subadults in the mesopelagic zone, with no adult specimens recorded (Guerra, González et al. 2014). It is expected that this species undergoes ontogenic migration, where indvividuals migrate to deeper waters as they develop, with adults inhabiting areas that match the foraging zones of sperm whales. Despite being present in 80% of the sperm whale faeces, *O. carriboea* was not found in n the mesopelagic zone, suggesting that it may not be among the most common and abundant species. Conversely, some known sperm whale prey found in the water samples, such as *T. pavo* and *Haliphron atlanticus* (Clarke, Martins et al. 1993), were not detected in the faeces.

Although cephalopods were the predominant prey in sperm whale samples, deep-sea fish were also found, with *C. couesii* being the main fish species. This species has never been recorded in the stomach contents of sperm whales from the Azores (Clarke 1956), and the only record in the literature was in a specimen from Arctic waters (Kharin 2006). *C.couesii* is a small (maximum length 7.5 cm) bathypelagic species commonly found in the lower mesopelagic zone in the Atlantic (500-1200m) (Pietsch 2002). Nevertheless, it was not detected in the water samples of this study. Taken together, it seems likely that this prey represents secondary consumption; in the absence of other fish DNA in the samples, it had no molecular competition and overamplified.

### Towards cetacean conservation by monitoring fish and cephalopod prey communities using multi-source DNA

Our study demonstrates the potential of combining environmental and fecal DNA to provide crucial insights for cetacean conservation. Environmental DNA data captured the known DVMs of mesopelagic fish species and the vertical distribution pattern of fish and cephalopod diversity, with most species found in the mesopelagic zone (Smith and Brown 2002, Rosa, Dierssen et al. 2008). Meanwhile, faecal DNA data provided a description of fish and cephalopods as prey of cetaceans, offering a level of detail that cannot be achieved through morphological or isotope analysis. Additionally, the combined analyses of these data sources have allowed us to describe preferences in prey consumption based on fish and cephalopod species availability. This information assists in identifying key prey for rorquals and sperm whales to improve the understanding of cetacean ecology and predict potential indirect risks associated with prey distribution and abundance. Additionally, the presence of DNA sequences that could only be classified to family level reveal the presence unknown diversity, which is documented for deep-sea environments due to the difficulty to study this remote and hostile ecosystem (Paulus 2021). These knowledge gaps complicate predictions of community responses to global changes (Jennings and Brander 2010, Poloczanska, Burrows et al. 2016) and call for addressing them for ensuring ecosystem-scale conservation. Overall, this study lays the groundwork for future research applying DNA analyses for studying cetacean diets, especially those species for which food web interaction information is lacking. This innovative approach will enable researchers to gain a deeper understanding of ecological and dietary interactions within marine ecosystems, allowing to advance towards ecosystem-based management, significantly contributing to the global conservation of marine biodiversity.

## Acknowledgments

Authors are grateful to the crew of the R/V OceanXplorer for helping with the water sampling and to the people involved in the surveys where faeces were collected. Authors are thankful to AMBAR Elkartea for collecting cetacean samples from stranded individuals, to the specimen biobank of the University of the Basque Country in the Plentzia Marine Station (PiE), specially to Xabier Lekube, for sharing samples for the development of the blocking primer used in this study, and to the University of Valencia, especially to Natalia Fraija, for sharing a sperm whale tissue sample. This work has been funded by the Portuguese Science & Technology Foundation (FCT) - project WATCH IT (Acores-01-0145-FEDER-000057), the Azorean Science & Technology Fund (FRCT) – project FCT Exploratory (IF/00943/2013/CP1199/CT0001), the European Union’s Horizon 2020 research and innovation program – project SUMMER (grant agreement No. 817806) co-funded by COMPETE, QREN, POPH, ESF, PO AZORES 2020 and Portuguese Ministry for Science and Education, and the Department of Economic Development and Infrastructure (Basque Government) - project GENGES. M.A.S and R.P. were supported by PO AZORES 2020 (MarAz Fund 01-0145-FEDER-000140) and project OCEAN (Horizon, GA 101076983), I.C. by FCT- IP Project UIDP/05634/2020, and C.O. by EUROPAMS (Biodiversa+, GA 101052342), and C.C by a predoctoral grant from the Department of Education (Basque Government).

## Author contributions

C.C., M.A.S. and N.R-E. conceived the study. R.P., I.C., M.A.S. and C.O. collected the samples. I.M.

C.C. and L.G.A. performed the laboratory work. C.C. and O.C. developed the code. C.C. wrote the first draft of the manuscript and M.A.S. and N.R-E. critically contributed with revisions. All authors gave the final approval for publication.

