## Appendix for "Contribution of mesopelagic fish and cephalopods to the diet of rorquals (*Balaenoptera spp*) and sperm whales (*Physeter macrocephalus*) beyond their feeding grounds"

**Supplementary material**

**Appendix 1: Supplementary Figures and Tables**

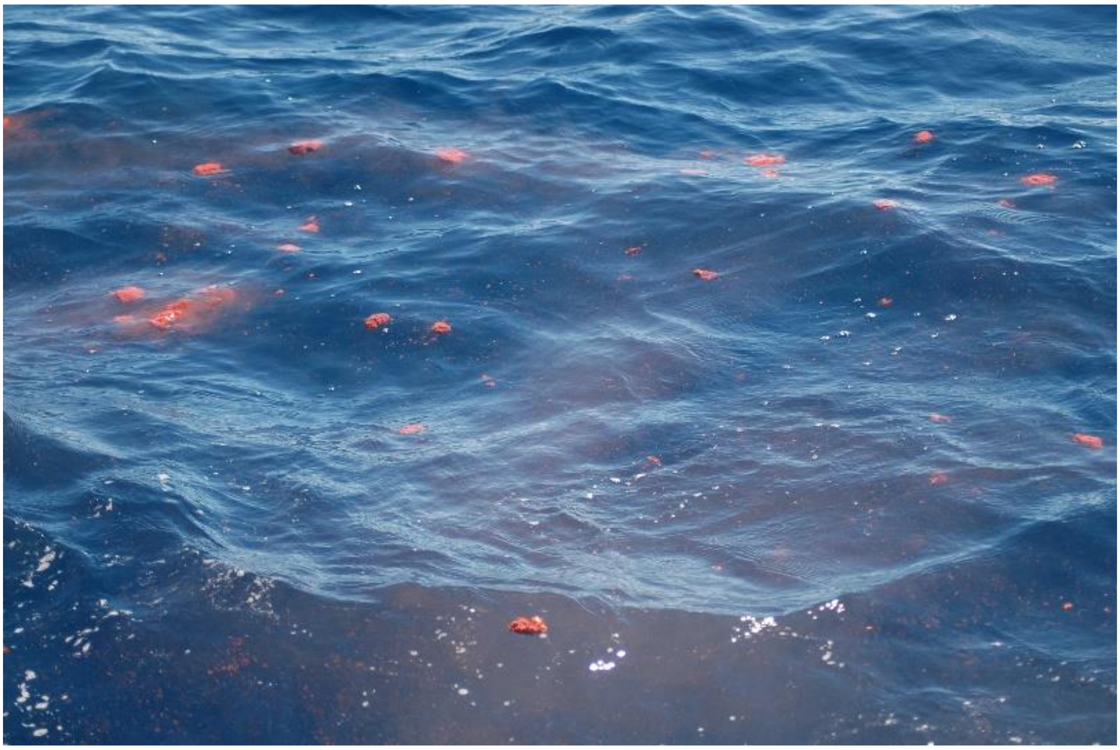

**Figure S1.** Baleen whale faeces floating in the water before collection. Samples were taken from the core of the biggest chunks.

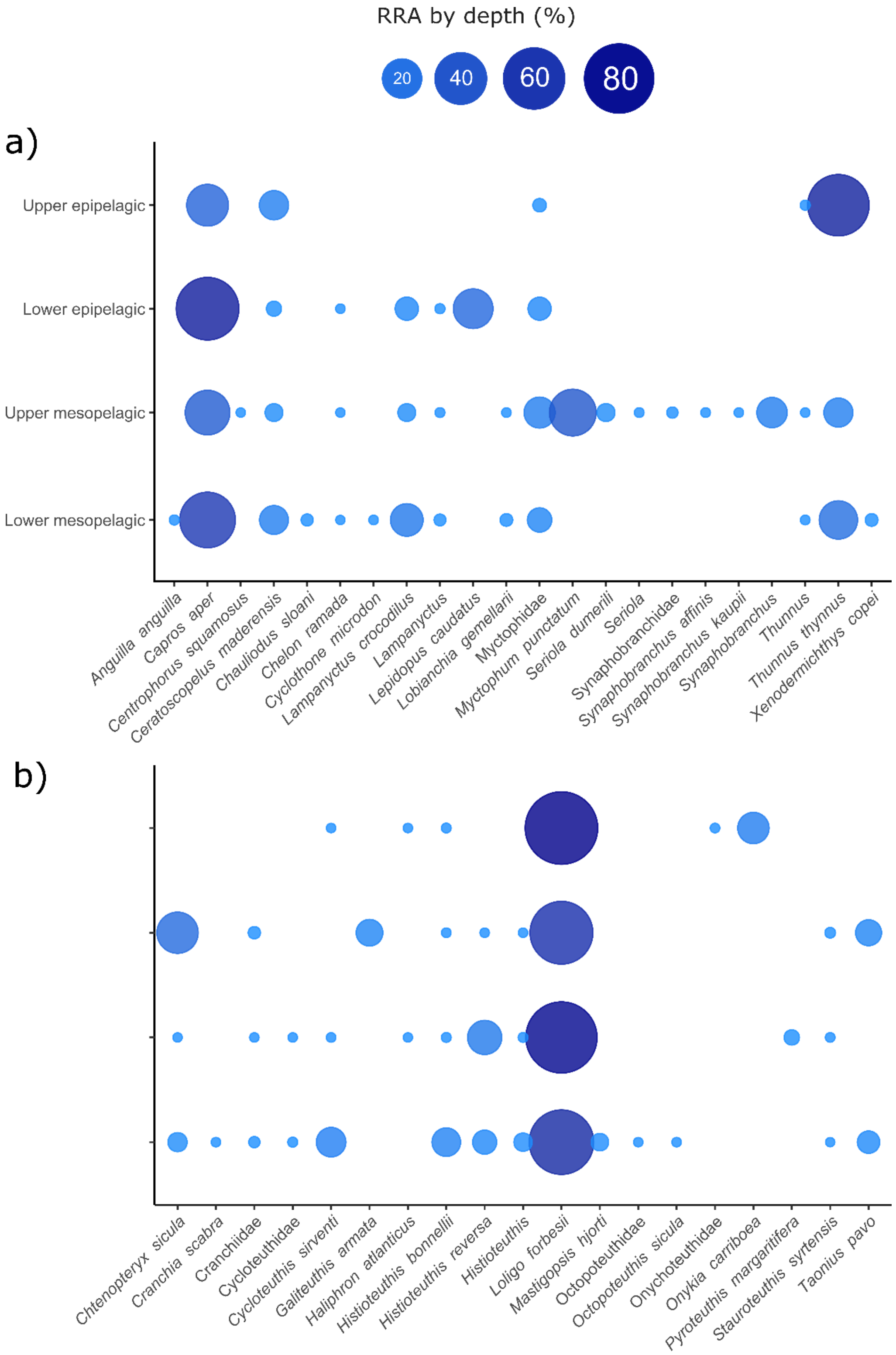


**Figure S2.** Bubble plots represent the vertical distribution of **a)** fish and **b)** cephalopod phylotypes detected in water samples, where the size of the bubbles indicates the relative read abundance (RRA, in %) of each species within a given depth range.

**Table S1.** Metadata of faecal samples, including the species, sampling location and date.

| Sample | Species | Location | Date |
|---|---|---|---|
| BBO_36_2 | *Balaenoptera borealis* | N Faial | 29/04/2014 |
| BMU_18 | *Balaeonoptera musculus* | S Faial | 14/04/2014 |
| BMU_25 | *Balaeonoptera musculus* | S Pico | 30/05/2014 |
| BMU_29 | *Balaeonoptera musculus* | SW Faial | 03/05/2016 |
| BMU_31 | *Balaeonoptera musculus* | S Pico | 13/05/2016 |
| BPH_03 | *Baaleonoptera physalis* | S Pico | 23/05/2011 |
| BPH_104 | *Balaeonoptera physalis* | N Faial | 29/05/2015 |
| BPH_105 | *Balaeonoptera physalis* | N Faial | 29/05/2015 |
| BPH_106 | *Balaeonoptera physalis* | N Faial | 29/05/2015 |
| BPH_107 | *Balaeonoptera physalis* | N Faial | 01/06/2015 |
| BPH_108 | *Balaeonoptera physalis* | N Faial | 01/06/2015 |
| BPH_109 | *Balaeonoptera physalis* | N Faial | 01/06/2015 |
| BPH_111 | *Balaeonoptera physalis* | N Faial | 08/06/2015 |
| BPH_121 | *Balaeonoptera physalis* | N Faial | 18/06/2015 |
| BPH_125_2 | *Balaeonoptera physalis* | S Faial | 23/06/2015 |
| BPH_146 | *Balaeonoptera physalis* | S Pico | 29/05/2017 |
| BPH_44 | *Balaeonoptera physalis* | S Pico | 23/05/2014 |
| BPH_48 | *Balaeonoptera physalis* | S Pico | 28/05/2014 |
| BPH_49 | *Balaeonoptera physalis* | S Pico | 30/05/2014 |
| BPH_53 | *Balaeonoptera physalis* | S Pico | 02/06/2014 |
| BPH_55 | *Balaeonoptera physalis* | S Faial | 19/06/2014 |
| BPH_57 | *Balaeonoptera physalis* | S Pico | 20/06/2014 |
| BPH_58 | *Balaeonoptera physalis* | S Pico | 20/06/2014 |
| BPH_63_2 | *Balaeonoptera physalis* | S Faial | 01/07/2014 |
| BPH_69 | *Balaeonoptera physalis* | N Faial | 04/05/2015 |
| PMA_185 | *Physeter macrocephalus* | S Pico | 11/08/2014 |
| PMA_271 | *Physeter macrocephalus* | Pico-S Jorge | 12/08/2020 |
| PMA_272 | *Physeter macrocephalus* | Pico-S Jorge | 12/08/2020 |
| PMA_274 | *Physeter macrocephalus* | Pico-S Jorge | 12/08/2020 |
| PMA_275 | *Physeter macrocephalus* | Pico-S Jorge | 12/08/2020 |
| PMA_282 | *Physeter macrocephalus* | S Pico | 28/07/2021 |
| PMA_285 | *Physeter macrocephalus* | S Pico | 03/08/2021 |
| PMA_287 | *Physeter macrocephalus* | S Pico | 03/08/2021 |
| PMA_288 | *Physeter macrocephalus* | S Pico | 09/08/2021 |

**Table S2.** Metadata of vertical profiles, including sampling depth, location, date and moment.

| Sample | Depth (m) | Location | Date | Sampling moment |
|---|---|---|---|---|
| AZO01 | 1200 | NW São Jorge | 13/06/2021 | night |
| AZO02 | 1001 | NW São Jorge | 13/06/2021 | night |
| AZO03 | 800 | NW São Jorge | 13/06/2021 | night |
| AZO04 | 601 | NW São Jorge | 13/06/2021 | night |
| AZO05 | 402 | NW São Jorge | 13/06/2021 | night |
| AZO06 | 200 | NW São Jorge | 13/06/2021 | night |
| AZO07 | 60 | NW São Jorge | 13/06/2021 | night |
| AZO08 | 10 | NW São Jorge | 13/06/2021 | night |
| AZO09 | 1200 | WNW Graciosa | 14/06/2021 | day |
| AZO10 | 1000 | WNW Graciosa | 14/06/2021 | day |
| AZO11 | 800 | WNW Graciosa | 14/06/2021 | day |
| AZO12 | 601 | WNW Graciosa | 14/06/2021 | day |
| AZO13 | 398 | WNW Graciosa | 14/06/2021 | day |
| AZO14 | 200 | WNW Graciosa | 14/06/2021 | day |
| AZO15 | 62 | WNW Graciosa | 14/06/2021 | day |
| AZO16 | 10 | WNW Graciosa | 14/06/2021 | day |
| AZO17 | 800 | SW Graciosa | 15/06/2021 | day |
| AZO18 | 600 | SW Graciosa | 15/06/2021 | day |
| AZO19 | 350 | SW Graciosa | 15/06/2021 | day |
| AZO20 | 60 | SW Graciosa | 15/06/2021 | day |
| AZO21 | 10 | SW Graciosa | 15/06/2021 | day |
| AZO22 | 792 | SW Graciosa | 16/06/2021 | night |
| AZO23 | 601 | SW Graciosa | 16/06/2021 | night |
| AZO24 | 349 | SW Graciosa | 16/06/2021 | night |
| AZO25 | 61 | SW Graciosa | 16/06/2021 | night |
| AZO26 | 11 | SW Graciosa | 16/06/2021 | night |
| AZO27 | 990 | N São Jorge | 18/06/2021 | day |
| AZO28 | 600 | N São Jorge | 18/06/2021 | day |
| AZO29 | 400 | N São Jorge | 18/06/2021 | day |
| AZO30 | 199 | N São Jorge | 18/06/2021 | day |
| AZO31 | 60 | N São Jorge | 18/06/2021 | day |
| AZO32 | 996 | NW São Jorge | 18/06/2021 | night |
| AZO33 | 600 | NW São Jorge | 18/06/2021 | night |
| AZO34 | 399 | NW São Jorge | 18/06/2021 | night |
| AZO35 | 200 | NW São Jorge | 18/06/2021 | night |
| AZO36 | 60 | NW São Jorge | 18/06/2021 | night |
| AZO38 | 800 | W Graciosa | 21/06/2021 | day |
| AZO39 | 630 | W Graciosa | 21/06/2021 | day |
| AZO40 | 430 | W Graciosa | 21/06/2021 | day |
| AZO41 | 100 | W Graciosa | 21/06/2021 | day |
| AZO42 | 10 | W Graciosa | 21/06/2021 | day |
| AZO43 | 800 | W Graciosa | 21/06/2021 | night |
| AZO44 | 630 | W Graciosa | 21/06/2021 | night |
| AZO45 | 430 | W Graciosa | 21/06/2021 | night |
| AZO46 | 100 | W Graciosa | 21/06/2021 | night |
| AZO47 | 10 | W Graciosa | 21/06/2021 | night |
| AZO48 | 980 | N Graciosa | 22/06/2021 | day |

| | | | | |
|---|---|---|---|---|
| AZO49 | 500 | N Graciosa | 22/06/2021 | day |
| AZO50 | 300 | N Graciosa | 22/06/2021 | day |
| AZO51 | 60 | N Graciosa | 22/06/2021 | day |
| AZO52 | 10 | N Graciosa | 22/06/2021 | day |
| AZO53 | 973 | N Graciosa | 24/06/2021 | night |
| AZO54 | 500 | N Graciosa | 24/06/2021 | night |
| AZO55 | 300 | N Graciosa | 24/06/2021 | night |
| AZO56 | 59 | N Graciosa | 24/06/2021 | night |
| AZO57 | 10 | N Graciosa | 24/06/2021 | night |
| AZO58 | 660 | S Faial-Pico | 26/06/2021 | night |
| AZO59 | 560 | S Faial-Pico | 26/06/2021 | night |
| AZO60 | 300 | S Faial-Pico | 26/06/2021 | night |
| AZO61 | 100 | S Faial-Pico | 26/06/2021 | night |
| AZO62 | 10 | S Faial-Pico | 26/06/2021 | night |
| AZO63 | 663 | S Faial-Pico | 27/06/2021 | day |
| AZO64 | 560 | S Faial-Pico | 27/06/2021 | day |
| AZO65 | 300 | S Faial-Pico | 27/06/2021 | day |
| AZO66 | 100 | S Faial-Pico | 27/06/2021 | day |
| AZO67 | 10 | S Faial-Pico | 27/06/2021 | day |

**Table S3.** For water samples, number of fish reads obtained and percentage of reads assigned to family, genus or species phylotypes.

| | | Assignation level | | |
|---|---|---|---|---|
| Depth range category | Fish reads | Species | Genus | Family |
| Upper epipelagic | 1,136,471 | 99.555 | 0.018 | 0.427 |
| Lower epipelagic | 331,833 | 95.361 | 0.017 | 4.622 |
| Upper mesopelagic | 765,190 | 77.886 | 10.648 | 11.467 |
| Lower mesopelagic | 819,765 | 94.351 | 0.168 | 5.481 |

**Table S4.** For water samples, number of cephalopod reads obtained, and percentage of reads assigned to family, genus or species phylotypes.

| | | Assignation level | | |
|---|---|---|---|---|
| Depth range category | Cephalopod reads | Species | Genus | Family |
| Upper epipelagic | 1,773,833 | 99.997 | 0 | 0.003 |
| Lower epipelagic | 1,584,099 | 99.794 | 0.001 | 0.205 |
| Upper mesopelagic | 4,642,465 | 99.992 | 0.007 | 0.001 |
| Lower mesopelagic | 10,577,486 | 98.089 | 1.813 | 0.098 |

**Table S5.** For faecal samples, number of fish reads obtained, and percentage of reads assigned to family, genus or species phylotypes.

| | | Assignation level | | |
|---|---|---|---|---|
| **Sample** | **Fish reads** | **Species** | **Genus** | **Family** |
| TELBBO_36_2 | 10,149 | 99.882 | 0.118 | 0 |
| TELBMU_18 | 31,083 | 0 | 0 | 100 |
| TELBMU_25_2 | 20,863 | 100 | 0 | 0 |
| TELBPH_03 | 61,575 | 100 | 0 | 0 |
| TELBPH_104 | 7,137 | 100 | 0 | 0 |
| TELBPH_105 | 10,149 | 100 | 0 | 0 |
| TELBPH_107 | 3,396 | 100 | 0 | 0 |
| TELBPH_108 | 11,286 | 100 | 0 | 0 |
| TELBPH_109 | 3,650 | 100 | 0 | 0 |
| TELBPH_121 | 59,494 | 99.955 | 0.024 | 0.022 |
| TELBPH_125_2 | 25,077 | 99.960 | 0 | 0.040 |
| TELBPH_146 | 5,506 | 100 | 0 | 0 |
| TELBPH_44 | 84,567 | 93.391 | 0 | 6.609 |
| TELBPH_48 | 65,819 | 99.982 | 0.018 | 0 |
| TELBPH_49 | 205,967 | 99.972 | 0.028 | 0 |
| TELBPH_57 | 122,481 | 99.973 | 0.027 | 0 |
| TELBPH_58 | 16,272 | 100 | 0 | 0 |
| TELBPH_63_2 | 161 | 100 | 0 | 0 |
| TELBPH_69 | 70,002 | 99.974 | 0.026 | 0 |
| TELPMA_185 | 547 | 100 | 0 | 0 |
| TELPMA_271 | 9,136 | 100 | 0 | 0 |
| TELPMA_272 | 2,461 | 100 | 0 | 0 |
| TELPMA_275 | 234 | 100 | 0 | 0 |
| TELPMA_282 | 396 | 100 | 0 | 0 |
| TELPMA_285 | 64,547 | 3.525 | 0 | 96.475 |
| TELPMA_288 | 1,826 | 100 | 0 | 0 |

**Table S6.** For faecal samples, number of cephalopod reads obtained, and percentage of reads assigned to family, genus or species phylotypes.

| | | Assignation level | | |
|---|---|---|---|---|
| **Sample** | **Cephalopod reads** | **Species** | **Genus** | **Family** |
| CEPBBO_36_2 | 653 | 88.668 | 6.126 | 5.207 |
| CEPBMU_18 | 122 | 100 | 0 | 0 |
| CEPBMU_25 | 30 | 33.333 | 0 | 66.667 |
| CEPBMU_29 | 325,709 | 99.905 | 0 | 0.095 |
| CEPBMU_31 | 68 | 100 | 0 | 0 |
| CEPBPH_03 | 28,329 | 100 | 0 | 0 |
| CEPBPH_104 | 129 | 71.318 | 0 | 28.682 |
| CEPBPH_105 | 270,763 | 99.916 | 0.040 | 0.044 |
| CEPBPH_107 | 769 | 98.440 | 1.560 | 0 |
| CEPBPH_108 | 16 | 100 | 0 | 0 |
| CEPBPH_109 | 9,171 | 100 | 0 | 0 |
| CEPBPH_111 | 64 | 100 | 0 | 0 |
| CEPBPH_121 | 33 | 100 | 0 | 0 |
| CEPBPH_125_2 | 14 | 100 | 0 | 0 |
| CEPBPH_44 | 200 | 100 | 0 | 0 |
| CEPBPH_48 | 8,640 | 100 | 0 | 0 |
| CEPBPH_49 | 3,588 | 100 | 0 | 0 |
| CEPBPH_55 | 85 | 100 | 0 | 0 |
| CEPBPH_57 | 60 | 100 | 0 | 0 |
| CEPBPH_58 | 41 | 100 | 0 | 0 |
| CEPBPH_63_2 | 137 | 82.482 | 0 | 17.518 |
| CEPBPH_69 | 404,098 | 100 | 0 | 0 |
| CEPPMA_185 | 1,184,698 | 96.759 | 3.194 | 0.047 |
| CEPPMA_271 | 3,954,457 | 94.249 | 5.713 | 0.038 |
| CEPPMA_272 | 220,141 | 95.647 | 4.353 | 0 |
| CEPPMA_274 | 105,355 | 90.755 | 8.641 | 0.604 |
| CEPPMA_275 | 3,159,807 | 96.783 | 3.049 | 0.168 |
| CEPPMA_282 | 1,940,653 | 56.014 | 43.985 | 0.002 |
| CEPPMA_285 | 885,127 | 98.569 | 1.352 | 0.079 |
| CEPPMA_287 | 1,632,633 | 97.405 | 2.594 | 0.001 |
| CEPPMA_288 | 4,730,077 | 98.941 | 1.059 | 0.000 |

**Table S7.** Fish phylotype table, as separated excel file.

**Table S8.** Cephalopod phylotype table, as separated excel file.

<u>**Appendix 2: Supplementary Materials and Methods**</u>

<u>Blocking primer generation and testing</u>

Faecal samples have high amounts of good quality predator DNA compared to degraded DNA of preys after digestion, which can lead to over-sequencing of predator DNA if co-amplified (Günther, Fromentin et al. 2021). To avoid nontarget amplification a blocking primer for suppressing whale DNA amplification was designed for the *teleo* forward primer following recommendations described in Vestheim, Deagle et al. (2011).

First, all available 12S sequences (including mitogenomes) belonging to the whale species included in this study (*Balaenoptera physalus, Balaenoptera musculus, Balaenoptera borealis* and *Physeter macrocephalus*) were downloaded from GenBank (https://www.ncbi.nlm.nih.gov/nucleotide/). Sequences were aligned with MAFFT (Katoh, Misawa et al. 2002) and a consensus predator sequence was created using BioEdit (Hall 1999), taken into consideration the Atlantic haplotypes of rorquals. The 12S region including the *teleo* amplicon and primers was trimmed. Second, a representative group of fish prey species were identified from bibliography (Watkins and Schevill 1979, Kawakami 1980, Clarke, Martins et al. 1993, Horwood 2009, Aguilar and García-Vernet 2018, Carwardine 2019) and 12S available sequences were retrieved from GenBank, aligned and *teleo* region trimmed as described above. Third, the predator consensus sequence, prey sequences and the *teleo* forward and reverse primers were aligned (**Figure A**).

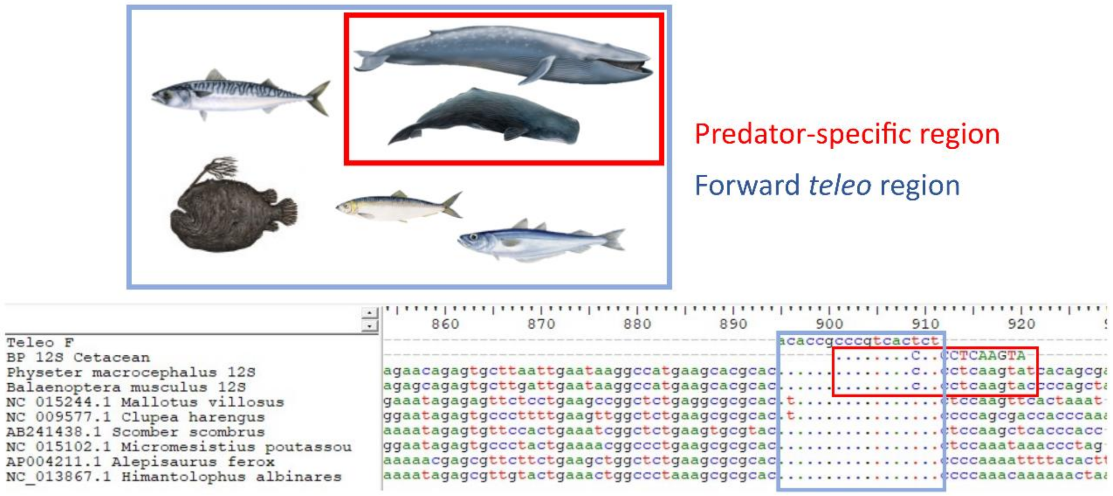


**Figure A.** Alignment of the predator and prey sequences with the *teleo* forward primer.

Entropy was calculated with BioEdit for each position to find hypervariable and conserved regions. A suitable area in the alignment for placement of the probe (i.e., unique to cetaceans meant to be blocked out) was selected overlapping the forward *teleo* primer. The melting temperature, hairpin formation and self-dimerisation (**Figure B**) of the probe were analysed with the OligoAnalyzer™ Tool (IntegratedDNATechnologies) to verify its feasibility. The probe was tested *in silico* using BLAST and resulted to have no complementarity with any marine species.

```
Delta G: -3.61 kcal/mole  Base Pairs: 2
5'                CCCGTCACCCTCCTCAAGTA
                      ||
3' ATGAACTCCTCCCACTGCCC
```

```
Delta G: -1.6 kcal/mole  Base Pairs: 2
5' CCCGTCACCCTCCTCAAGTA
            ||      ::
3'        ATGAACTCCTCCCACTGCCC
```

```
Delta G: -1.6 kcal/mole  Base Pairs: 2
5' CCCGTCACCCTCCTCAAGTA
              : ||  :: :
3'           ATGAACTCCTCCCACTGCCC
```

```
Delta G: -1.34 kcal/mole  Base Pairs: 2
5'           CCCGTCACCCTCCTCAAGTA
                ||  ::
3' ATGAACTCCTCCCACTGCCC
```

**Structures**

| structure | Image | ΔG (kcal.mole$^{-1}$) | $T_m$ (°C) | ΔH (kcal.mole$^{-1}$) | ΔS (cal.K$^{-1}$mole$^{-1}$) |
|---|---|---|---|---|---|
| 1 | 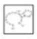 | 0.54 | 16.9 | -19.2 | -66.2 |
| 2 | 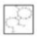 | 1.48 | -18.4 | -8.7 | -34.15 |

**Figure B.** Hairpin formation and self-dimerisation test of the bocking primer.

To test the probe *in vitro*, predator and prey (several Atlantic fish species) DNA extractions were performed from tissue using the Wizard Genomic DNA Purification Kit (Promega) following manufacturer's instructions for mouse tail extraction. Amplifications were preformed using qPCR. Different proportions of the blocking primer and the *teleo* primers were tested in know predator-prey mixes. The probe resulted in a 20 nucleotide sequence and includes a C3 Spacer (3 hydrocarbons) in the 3´ that blocks the annealing (Vestheim and Jarman 2008). For the PCR step, 0.3 µl of blocking primer at [2µM] were included to each sample to the amplification mix.

- BP_12S_Cetacea (5´-3´): CCCGTCACCCTCCTCAAGTA/SpC3

Specifications and characteristics based on Vestheim et al. (2011):

- Tm = 58.4°C (Slightly higher than *teleo* Tm, 55°C)
- Length: 20 bp (< 25bp)
- No more than 3 G/C content at the 3' end
- No long runs of single bases (> 3bp)
- No more than 4 dinucleotide repeats

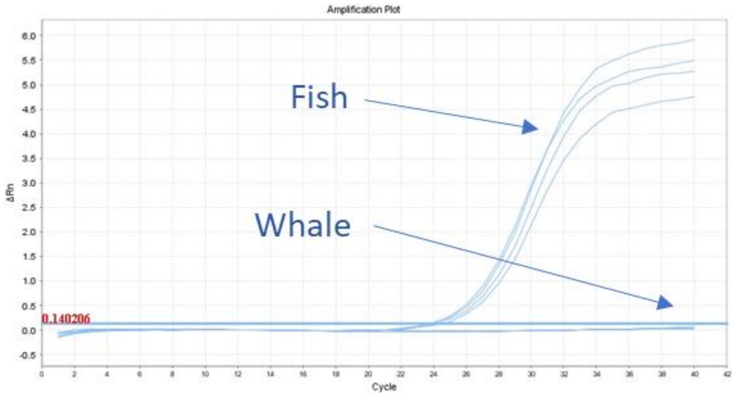


**Figure C.** Amplification plot of tissue samples of fish (mock communities designed in-house) and whales (*B. physalus, P. macrocephalus*) adding designed blocking primers.
